# Heart evoked brain oscillatory networks and its interruption in early stages of Alzheimer’s disease

**DOI:** 10.1101/407361

**Authors:** Nazareth Castellanos, Gustavo G. Diez, Ernesto Pereda, María Eugenia López, Ricardo Bruña, Myriam G. Bartolomé, Fernando Maestú

## Abstract

Understanding how the heart influences brain dynamics will suppose a deep change for the neuroscience, psychology and medicine. A mainstay questions is the heart modulation of resting state brain networks and its relation with both the cardiac dynamics and the cognitive status. We evaluated the heart evoked basal networks for controls and two groups of mild cognitive impairment patients, stable and progressive to Alzheimer’s disease without cardiovascular alteration symptoms. Our results in controls show that a healthy cognitive performance correlates with the heart modulation of brain dynamics in areas of the default mode network, and that the heart influence on brain networks varies along the cardiac cycle and the spectral band. However, the cognitive deficit produced by dementia correlates with the lack of heart modulation on brain activity. The heart influence on brain networks is disrupted in patients by producing hypersynchronization, accompanied by decreased cardiac complexity. We designed a surrogate and predictive procedure based on machine learning to compare the heart evoked results with the neural activity no locked to heartbeats. Based on our longitudinal data, we conclude that the prediction to progression to Alzheimer’s disease is higher when considering the heart - brain interaction than when taking into account only the brain dynamics. We can conclude that brain networks in control subjects were more responsive to the heart cycle, allowing a wealthier, more complex pattern of oscillations. Our results highlight the role of heart in cognitive neuroscience by showing that basal brain networks are modulated by the cardiac dynamics.

## Introduction

The brain-body-mind interaction is a central issue in the philosophy of mind, although historically it has not been related to the physical body. The progress in physiological techniques and the advanced knowledge of biological mechanisms are reviving this fundamental philosophical question, which can dramatically change our basic concepts in medicine, opening new perspectives for intervention. A recent ground-breaking hypothesis in cognitive neuroscience grants an active role to heart activity in influencing neuronal dynamics. The study of the interaction between brain and heart represents a fundamental step in human functional neuroimaging, involving functional magnetic resonance imaging (fMRI), electroencephalography (EEG) and magnetoencephalography (MEG) studies, and opens the door to conceive the mind as a dynamically embodied system (Varela et al., 1992). Pioneering works (Park et al., 2014, Gray et al., 2009, Critchley et al., 2004) suggest that the response of the brain to the heart pulse determines the outcome of cognitive tasks, which some authors have interpreted (Park et al., 2016) as a measure of the subjective feeling of the self in the perceptual moment, favouring a successful detection of a visual neutral stimulus. The heart-evoked neuronal response (HER) can be detected in different neocortical areas (Kern et al., 2003, Gray et al., 2007) such as the insula (Craig, 2002), the anterior cingulate cortex (Amour and Ardell, 2004, Vogt and Derbyshire, 2009), the amygdala (Craig, 2002) and the somatosensory cortex (Kern et al., 2003). This activity is currently associated with interoceptive and empathic abilities (Montoya et al., 1993, Fukushima et al., 2011), but it was historically interpreted as the viscerosensory processing in the brain (Critchley et al., 2004).

The role of heart would deeply change our conception of information processing but also importantly will open new doors in the clinical interventions of diseases considered purely of brain origin. This is the case of Alzheimer disease. Previous studies in dementia have associated heart and brain disease (Qiu and Fratiglioni, 2015), indicating that the most common form of dementia courses with cerebrovascular lesions, which can be one of the causes of neurodegeneration. Subclinical cardiovascular alterations are related to cognitive decline and dementia by causing cerebral hypoxia and silent brain injuries. In fact, dementia and cardiovascular diseases (CVD) share most of the vascular risk factors which, when accumulated during adulthood, substantially increase the risk of dementia (Sposato et al., 2017). Instead, cardiovascular health improves cognitive performance and reduces the risk of cognitive impairment (Crichton et al., 2014, Thacker, 2014). Although the cortical representation of the heartbeat includes areas of the default mode network (DMN), whose activity is largely involved in different stages of dementia (Babo-Rebelo et al., 2016), there is no evidence of this heart-brain relation in resting state (RS).

The question of whether this cardiovascular damage could be one of the factors responsible for brain network malfunctioning is being very recently discussed (Chong et al., 2017). Here, we hypothesize that underlying these results is the fact that heart activity interfering with brain oscillatory networks.

## Materials and Methods

### Subjects

We enrolled 53 patients diagnosed with amnestic-Mild Cognitive Impairment (MCI) according to the National Institute on Aging-Alzheimer Association (NIA-AA) criteria (Albert et al., 2011). They also showed significant hippocampal atrophy, which was evaluated by an experienced radiologist. Additionally, we carried out a clinical 3-year follow-up of the MCI subjects with the aim to determine if they either remained as MCIs or fulfilled the criteria for probable Alzheimer’s Disease according to the NIA-AA (McKhann et al., 2011). Based on their clinical outcome, MCI participants were then split into two subgroups: the stable MCI group (sMCI; n= 26), and the progressive MCI group (pMCI; n= 27). A sample of 26 age-matched controls was also selected with the same gender distribution and educational level of the MCI patients. Control and MCI patients differed in the Mini-mental state (MMSE) examination score (p < 0.001). All participants were in good health and had no history of psychiatric or other neurological disorders. The local Ethics Committee approved the investigation.

We determined the APOE genotype the blood samples of the MCI subjects using standard methods (Hixson et al., 1990). Participants were then classified according to the presence or absence of the ε4 allele as APOE4 carriers (if they exhibited at least one allele 4) or non-carriers (if they did not show any allele 4).

### MEG acquisition

Three-minute MEG resting-state recordings were acquired using an Elekta Vectorview system with 306 sensors, inside a magnetically shielded room (Vacuumschmelze GmbH, Hanau, Germany). To determine the head position inside the MEG helmet, we digitalized the head with a Fastrack Polhemus and four coils attached to the forehead and mastoids. Signals were sampled at 1 kHz with an online filter of bandwidth 0.1–300 Hz. Maxfilter software (version 2.2, Elekta Neuromag) was used to remove external noise with the temporal extension of the signal space separation (tsss) method with movement compensation (Taulu and Simola, 2006). The number of trials per subject is given by the number of heartpulses, being statistically similar in controls and patients (664 trials for controls and 587 for DCL).

### Estimation of the cardiac magnetic activity in the brain

The standard pre-processing of MEEG (MEG or EEG) data entails the rejection of components originated in external brain sources, such as eyes, muscles or the heart, to obtain a clean signal of the magnetic field of the brain. However, we consider that the cardiac magnetic component arriving at the brain far from being an artefact influences neuronal activity even in resting state. To estimate such cardiac component we applied Independent Component Analysis (Anemüller et al., 2003) as implemented in Fieldtrip (Oostenveld et al., 2011) to MEG segments of 4 seconds length. We manually selected the components of interest for this study, namely the neuronal and cardiac components for each segment and subject. Those components of neuronal origin were used for the source reconstruction method, avoiding the mixture with influences external to the brain. We individually checked the topography of the cardiac components and the presence of heartbeat and T wave, excluding those subjects with doubtful morphology. With this procedure, we expand MEG as a technique to study the brain activity to the Magneto-encephalo-cardiography (MEKG) to explore the brain-heart connection. ICA has been proved to be a robust method for artifact detection and is nowadays a standard technique used in the preprocessing of MEEG data. In this study we estimate the cardiac magnetic dynamics from the raw data by means of ICA (MEKG) instead of using a direct electrocardiogram (EKG). The first advantage of this procedure is to correlate both the brain and heart biomagnetic activities since the standard EKG estimates only the electric component of the heart electromagnetic field. Furthermore, with MEKG we estimate the cardiac magnetic field as arrived at the brain. As recently pointed (Winston and Rees, 2014), the possible distortion of the cardiac electromagnetic field from the heart to the brain is nowadays unknown. In order to validate our procedure we estimate the correlation between the manually extracted components from ICA and the EKG for a subset of participants (N = 7). We obtain a 100% of coincidence in the time location of heart pulses (1000 Hz sampling rate) very important since we estimate heart evoked brain modulation considering the pulse as a trigger. In addition, ICA algorithm supposes that the propagation delays from the sources to the electrodes are negligible. Respect to the dynamics, the correlation between both time series is, on average, of 87%. In addition, direct interaction between heart and brain dynamics is estimated by means of wavelet coherence (tempo-spectral counterpart of the correlation) based on the time-frequency representation of signals and hence less sensible to time variations in the estimation of the time series.

Although the heart rate and its variability are of great interest also in cognition, they are not a measure of the heart continuous dynamics. To characterise cardiac activity, we propose to estimate the complexity of the heart dynamics using mutual information (Shannon, 1948, Niso et al., 2013). Mutual Information (MI) is based on the concept of Shannon entropy, which is defined as the average amount of information gained from a measurement that specifies one particular value. Self-MI is a measure of the information of the values of a signal that is present on its past values. Thus, the value of MI is the uncertainty contained in one signal. The correlation time can be interpreted as the inner correlation or memory of the signal, measured as the delay (lag) in which the signal decays 1/e and takes into account both linear and non-linear inner correlations. This delay is considered here as the parameter measuring the memory of the cardiac magnetic component (EKG). We termed it as the “cardiac complexity or flexibility or, contrarily, stiffness” and used it to compare the cardiac dynamics of controls and MCI patients. We segmented the EKG in trials starting 50 ms before the QRS peaks and lasting until 50 ms before the next QRS peak. We have 664 trials for controls and 587 for DCL.

### MEG source reconstruction and heart evoked brain dynamics

After artifact rejection and the separation of the heart component, the 4 seconds segments of purely neuronal activity were used to estimate the sources in the standard spectral bands. Source locations were defined in the subject’s space by using the cortical segmentation produced by Freesurfer with a regular mesh of points with 1 cm spacing. We used a single shell model (Nolte, 2003) to solve the forward model and source reconstruction was estimated with Linearly Constrained Minimum Variance Beamformer (Van Veen et al., 1997) for each spectral band. Individually, spatial filter’s coefficients were computed by averaging the covariance matrix over all trials. Dynamics (time series) per segment and source localisation were obtained from these coefficients when applied to individual trials. To avoid mixing MEG sensors with different sensitivities or resorting to scaling, only magnetometers were used for source reconstruction. Note, however, that gradiometer information is indirectly present as both magnetometers and gradiometers were included in the tsss filtering.

Functional connectivity is calculated by means of wavelet coherence, estimates the interaction between two sources in both time and frequency simultaneously (Castellanos et al., 2012; Malmierca et al., 2009). We segmented the time courses of the sources and functional connectivity in trials centred on the QRS peak, called heart-evoked responses (HER) or heart-evoked brain network. In each of these trials, we defined four windows of interest: two of them around the QRS and T peaks ([-50, 100] ms) and two after the wave decay in the interval [200, 400] ms after both peaks, where brain activity is free from electrical artifacts due to heart contractions (Gray et al., 2007, Dirlich et al, 1998).

### Statistical analysis and machine learning classification

We used a nonparametric Mann–Whitney tests to check for differences between controls and MCIs in HER and connectivity. Further, we corrected by multiple comparisons by using a nonparametric permutation approach (Maris and Oosteveld, 2007), as follows. First, the original values were 5000 times randomly assigned to the original groups (controls and MCIs) and a Mann–Whitney test was performed for each randomization. Then, the U-value of the original dataset was compared to the ones obtained with the randomised data. The final p-value was defined as the proportion of permutations with U-values higher than the one of the original data. In all the correlations we estimated the normalized Pearson correlation coefficient. We established a threshold of abs(R) > 0.5 in order to avoid statistical correlation with a small slope for its risky interpretation. Statistical p-valus is thresholed to p<0.01 for all correlations showed in this work.

Additionally, we used a multivariate machine learning algorithm, support vector machine (SVM) (Vapnick, 1995), to classify stable and progressive MCI. The training phase (where the classifier was trained using group-labeled data) learns from the wavelet coherence between brain sources and heart dynamics from both sMCI and pMCI, and the accuracy is estimated from the testing phase with unseen data to be classified. A linear kernel was used to represent the data, reduce computational cost and improve classification accuracy (i.e., the overall rate of correct classification). An enhanced recursive feature procedure (Chen and Jeong, 2007) was implemented to ensure that discrimination accuracy was not due to overfitting and to select the best predictive features (brain areas). Finally, we used leave-one-out cross-validation (Geisser, 1993) to test classification accuracy and whether the results were independent of the initial training data.

## Results

### Heart evoked neuronal activity

The heart-evoked response (HER) was lower in MCI subjects than in controls (p<0.001, corrected) in three areas of the left hemisphere (the frontal orbital cortex, the frontal pole and the insular cortex), in the time window 200 ms after the QRS peak and centred in the T wave (Figure 1A). Further, we analysed the correlation, in MCIs and controls, between the HER and the results of different cognitive tests. In the MCI group, we also studied the possible relationship between HER and the degree of dementia as assessed by the MMSE. In both cases, we used the Pearson correlation coefficient R (at the p<0.01 level, threshold abs(R) > 0.5). HER correlated with different cognitive measures in several areas of the DMN (see Fig. 1C for their localization and the table at the bottom of this figure for the tests and the values of R). Besides, the score of the MMSE test correlated with the decrease of HER (Fig. 1B) in the left supracalcarine (R = 0.53) and precuneus cortices (R = 0.5) in the QRS peak, and in the left superior parietal lobe (R = 0.51) and paracingulate gyrus posterior (R = 0.55) 200 ms after the QRS peak.

**Figure 1:**
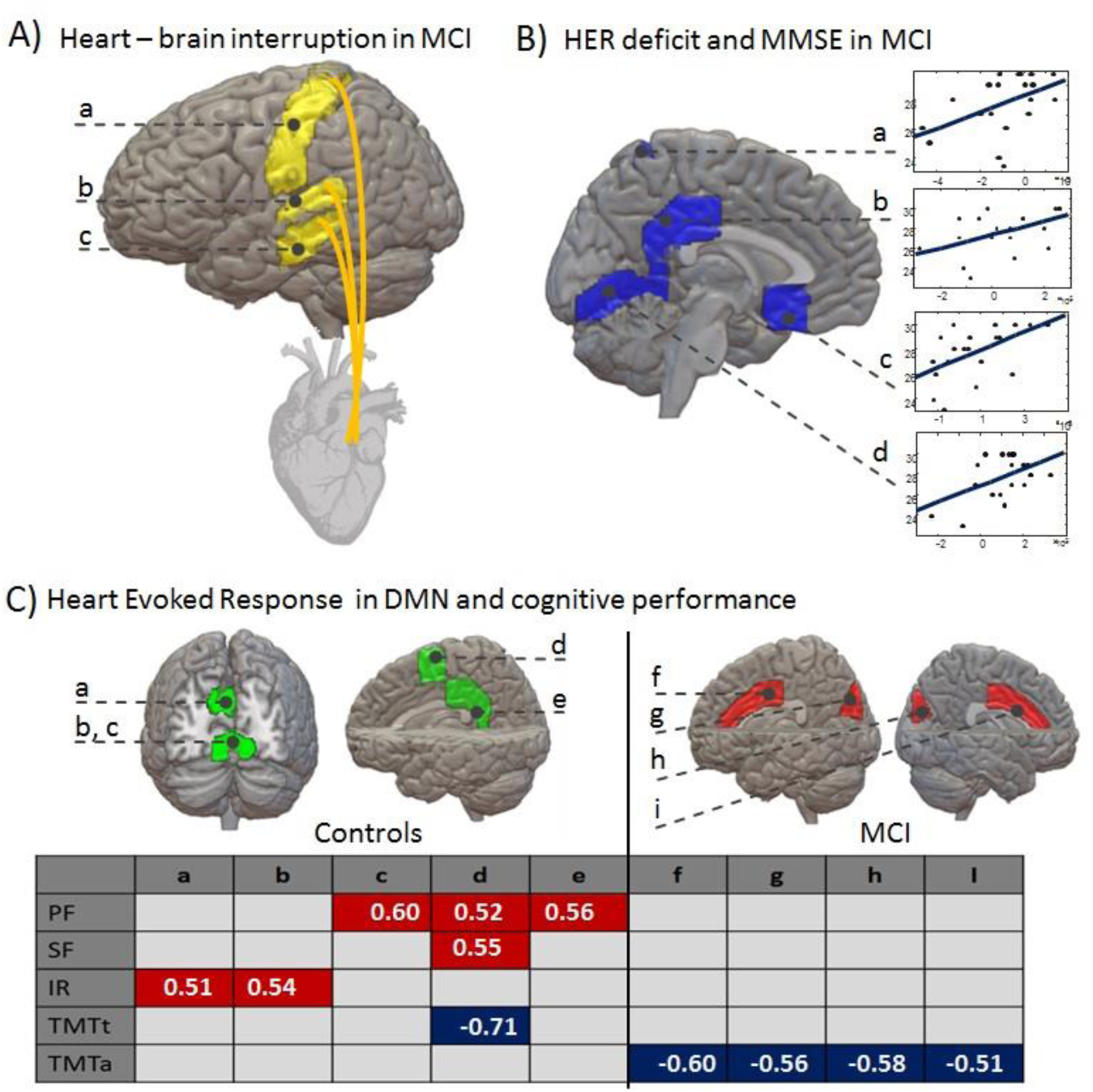
Heart evoked neuronal response and cognitive performance. A) Pattern of brain areas where heart modulation of brain activity is interrupted in MCI patients labelled as a) Left Frontal Pole, b) Left Insular cortex, and c) Left frontal orbital cortex. B) Correlation between HER with cognitive status as measured by MMSE in MCI patients. Areas labelled as: a) left precuneus cortex, b) left paracingulate gyrus posterior, c) left supracalcarine cortex, and d) left superior parietal lobe. C) Areas in the DMN presenting a significant correlation between HER and the cognitive test in control subjects (green areas) and MCI patients (red areas). Labels for brain areas: a) left parietal operculum cortex, b) left and c) right superior parietal lobe, d) right cuneal cortex, e) right precuneus, f) right supramarginal gyrus, g) Right Parietal Operculum cortex, h) Left Supramarginal gyrus and i) Left Parietal Operculum cortex (symmetrical areas not shown). Labels for cognitive tests: PF, phonetic fluency; SF, semantic fluency; IR, immediate recall; TMTt and TMTa, trail making test time and accuracy. Red color indicates positive correlation (R > 0.5) and blue color indicates negative correlation (R < −0.5).

### Heart modulation of brain functional connectivity

We assessed brain FC by using wavelet coherence, estimating the normalized weight of the functional interaction between two brain areas in both time and frequency simultaneously. To estimate heart-evoked FC, we took the QRS heartbeats as triggers and the previous segments of 100 ms as the basal FC state. Its statistical comparison allows inferring on the presence of a link between brain areas and its weight. A brain functional network is therefore defined by the weights and links between all brain areas. We analysed heart modulation of the brain network in the alpha band by looking at the time evolution of both its topology (number of significant links along time) and its global synchronization level (weight of these links). The influence of the heartbeat on network topology lasted longer in controls than in MCIs (Fig. 2A, left panel). Indeed, the impact of the heartbeat in brain network topology for MCI patients (red area) showed peaks at 50 and 200 ms, with the network going back to the pre-beat topology already after 240 ms. Instead, control subjects (blue area) showed a longer-lasting (625 ms) and more complex heart-evoked modulation of network topology. The heartbeat also modulated the overall synchronization level (Fig. 2A, right panel), showing a different behaviour in controls and MCIs: whereas it reduced global synchronization (especially in DMN areas) up to 650 ms after beating in control subjects, in MCIs the heartbeat induced hypersynchronization. Besides, this increase was stronger (in absolute value) in MCIs than in controls until 200 ms after the heartbeat. However, its modulation, regarding the temporal variability of its weight, was less accentuated than in controls. Brain networks in control subjects were more flexible and responsive to the heart cycle, allowing a wealthier, more complex pattern of oscillations. However, although the MCI network was hypersynchronized, its modulation during the cardiac cycle was reduced, which clearly suggests that the brain networks of MCI patients are stiffer and less flexible as compared to those of healthy age-matched subjects.

**Figure 2:**
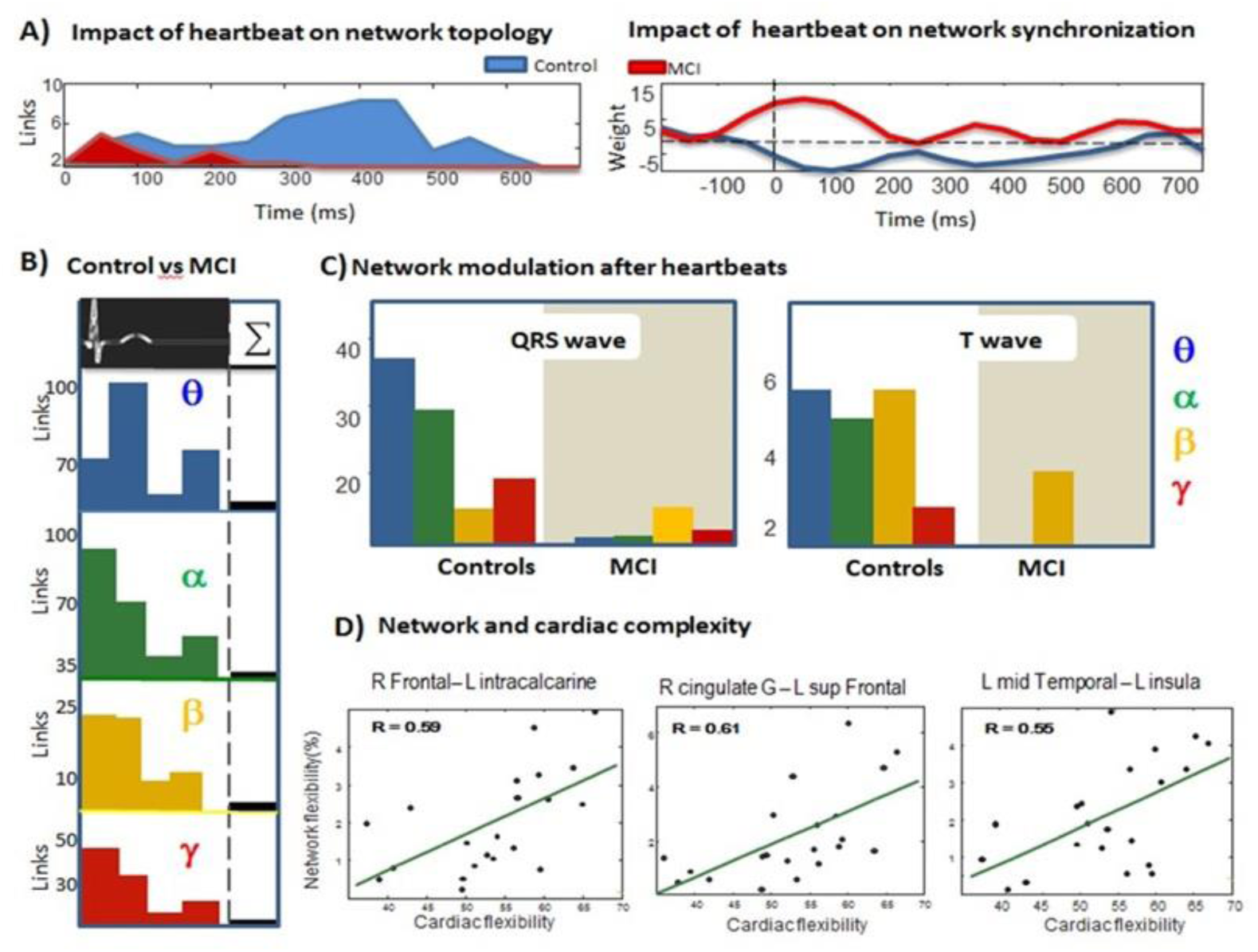
Heart modulation of brain network. A) Heart modulation of topology and global synchronization in alpha band brain network. The left panel shows the number of links statistically different from baseline for controls (blue area) and MCI subjects (red area). The right panel shows the percentage (%) of change (w.r.t. baseline) of network weight for controls (blue line) and MCI subjects (red line). Dashed line represents baseline reference. B) Global number of statistically different links between controls and MCI in spectral bands (colour code) along the heart cycle (4time windows from the QRS and T wave events [0, 150] and [200, 400] ms). The statistical topological differences between controls and MCI vary with time cycle. Black bars show the number of differential links without considering the cardiac events (i.e., brain activity not locked to heartbeats or non-reference resting state standard procedure). C) The global number of statistically different links after the QRS (left panel) and T-wave event (right panel) for controls (left) and MCI subjects (right) in spectral bands (colour code). Heartbeats in controls induce topological changes whereas in MCI subjects the network is stiffer. D) Correlation of network and cardiac flexibility in alpha based networks: the most the links change along the cardiac cycle in theta and alpha band, the greater cardiac flexibility.

Interestingly, between-group differences in the links within these brain networks were dynamic and depended on the instant of the cardiac cycle analysed. Indeed, they were highest in the QRS peak window and 200 ms later, especially in the alpha and beta bands (Fig. 2B). There is accumulated evidence that RS brain activity in MCI subjects is hypersynchronized in both bands (Bajo et al., 2010, Bajo et al., 2010b). Regionally, heart-evoked FC differences between controls and MCIs in the alpha band in the QRS window included a range of brain areas bilaterally. Namely, they were the temporal and occipital fusiform cortex, the temporal lobe, the insular cortex, the intracalcarine cortex, the precuneus and the frontal orbital cortex in the left hemisphere, and the frontal pole, medial and operculum cortex in the right one. In the second time window (200 ms after the QRS peak), the areas affected were both frontal poles, right occipital fusiform gyrus, left insular cortex and left intracalcarine cortex. Although we observed a hypersynchronization in alpha and beta bands in MCIs, the connection between left insular cortex and left frontal operculum cortex was missed in MCIs until 250 ms in both bands.

In order to test whether the results could be reproduced by using brain activity not locked to heartbeats we designed a surrogate procedure obtained by randomizing the temporal dependence of the cardiac pulse. Our hypothesis is that heart evoked FC differences between controls and patients are due to the heart modulation of brain FC. Our results are based on a statistical comparison of heart evoked brain networks in controls and patients considering the heartbeat as a trigger. To test whether same results can be inferred from brain activity alone (ignoring cardiac dynamics) we run the same analysis but randomly sampling the FC time series at intervals mimicking heartbeats (the standard procedure in RS studies). This comparison was repeated 10.000 times in order to estimate the rate of coincidences. The results of our simulations show that heart evoked FC are different from RS FC with less than 1/1000 coincidences (0.0007% of cases). RS is a non-reference condition where the beginning of the segments selected for analysis is typically chosen randomly. Clearly, such strategy results in a wastage of relevant information (black area in figure 2B), as compared to selecting the segments using the heartbeat as a trigger. In summary, heart-evoked FC differences between controls and MCI varies during the cardiac cycle, showing a greater modulation of pre-beat network topology in controls than in MCI subjects.

To study the impact of the R and T peaks on brain networks, we measured the dynamics of the links. As shown in Fig. 2C, the heart-evoked FC network of the MCI group was stiffer, changing less in all frequency bands than that of the control one. This lack of FC modulation correlated with cardiac complexity: the less the links changed along the cardiac cycle in theta and alpha band, the less the cardiac complexity (also called cardiac flexibility, Fig 2D). Interestingly, cardiac complexity correlated positively with FC between different brain areas in the theta band (Pearson correlation coefficient at p<0.01, corrected), with those between the left and right temporal, occipital fusiform gyrus (R = 0.62) and between the left temporal fusiform gyrus and right precuneus (R = 0.72). In the alpha band, with those between right frontal and left intracalcarine (R = 0.59), between right cingulate gyrus and left superior frontal (R = 0.61) and between the left middle temporal and left precuneus (R = 0.55).

### Heart – Brain synchronization predicts progression to AD and is influenced by APOE genotype

We used a multivariate machine learning technique (support vector machine, SVM) to test whether heart-brain synchronization is an accurate predictor of the progression of MCI subjects to AD. For this purpose, we estimated the accuracy of SVM to classify these subjects as sMCI or pMCI using heart-evoked brain synchronization and compared it with that obtained using brain synchronization alone. We used a recursive feature elimination algorithm to choose the most relevant time interval, spectral band and brain areas in both the heart-evoked and solely brain synchronization. The subset of brain areas performing better was later used in the classification. In this way, we not only prevented overfitting but also ensured that the number of features used for classification was the same in both cases. Figure 3 summarizes the outcome of the classifier: time interval [200-400 ms] after the heartbeat (T wave) in beta band and the areas right middle temporal gyrus, right frontal orbital cortex, left intracalcarine cortex, left precuneous cortex, right frontal operculum cortex, right supracalcarine cortex, right occipital fusiform gyrus, right intracalcarine cortex, right frontal medial cortex and left frontal orbital cortex. In all these regions, the heart-evoked brain synchronization was higher for the sMCI group. Notably, classification accuracy was greater when we used the heart-evoked synchronization values as compared to the situation when we classified the subjects using the brain-brain interaction only. Figure 4 shows prediction in classification accuracy for the progressive group when the brain-heart interaction is taken into account as compared to when only information from the brain-brain interaction is used. Finally, we analysed the brain heart synchronization evoked by the heartbeat according to the APOE genotype. We found that both MCI carriers of APOE4 showed higher synchronization in the beta band than MCI non-carriers in left frontal pole (p = 0.0027), left (p = 0.034) and right cingulate gyrus (p = 0.004), and right superior frontal gyrus (p = 0.002). APOE4 carriers were more likely to be hypersynchronized due to the accumulation of amyloid plaques. Furthermore, we observed that the sMCI carrier group was heart-evoked hypersynchronized as compared to pMCI APOE4 carriers in both the left and the right cingulate (p < 0.001).

**Figure 4:**
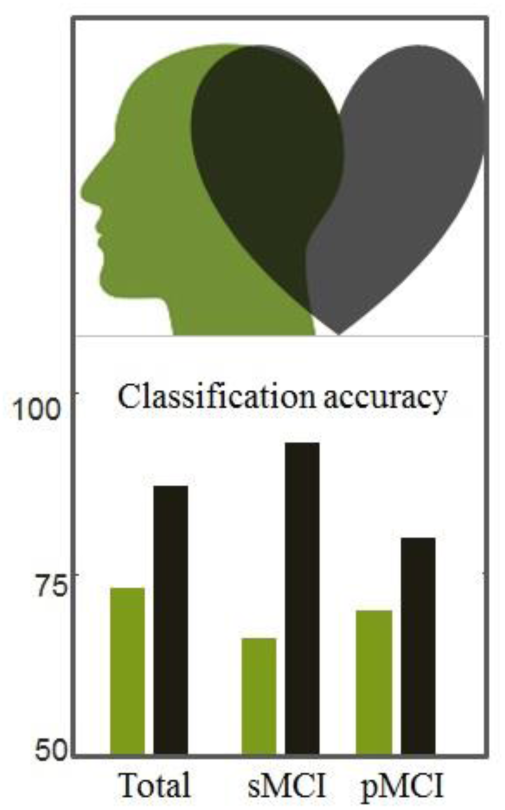
Improvement of Alzheimer disease prediction when considering both heart and brain synchronization as compared with purely brain. Classification accuracy (%) for stable and progressive MCI groups for the beta band in a time window [200, 400] ms after the heartbeat (as selected by a recursive feature selection).

## Discussion

### Clinical implications

We already know that the visceral status influences stimuli processing, requiring, therefore, the integration of sensorial information with autonomic control of the cardiovascular function. A question derived from this assumption is to what extent cardiac and brain malfunctioning appears simultaneously, and whether the former relates to the latter in diseases traditionally considered as a purely cerebral such as dementia (Qiu and Fratiglioni, 2015, Kisler et al., 2017). Several studies have reported that CVDs such as heart failure (Qui et al., 2005), hypertension (Thorin, 2015), atrial fibrillation (Kalantarian et al., 2016, Kwok et al., 2011) produce cognitive decline, either transitory or chronic. A recent rigorous and complete review (Qui and Fratiglioni, 2015) showed that the most common form of dementia courses with cerebrovascular lesions and neurodegeneration in elderly people. Post-mortem studies have also revealed that most cases of dementia involve brain lesions as macro and microinfarcts, white matter lesions, lacunar infarcts and cerebral microbleeds as well as brain degeneration such as neuritic plaques and neurofibrillary tangles. The reduction of at least five risk factors could prevent up to 3 million cases of Alzheimer disease (Barnes and Yaffe, 2011), as also supported by the fact that cardiovascular health improves cognitive performance and reduces the risk of cognitive impairment (Crichton et al, 2014).

The clinical implications of these findings are clearer once we consider the two factors studied here: the conversion from MCI to Alzheimer disease and the role of the APOE4 allele. We found that sMCI subjects showed higher synchronization than pMCIs in several brain regions. One plausible explanation is that sMCI subjects compensate cognitive deficits by increasing the rate of synchronization of their brain networks. Indeed, compensation is a common interpretation for the higher activation/synchronization in MCI subjects as compared to controls. However, such increased activity correlates with close-to-random organization of the MCI network and does not improve cognitive function. Furthermore, using a computational model, De Haan and colleagues (De Haan et al., 2017) demonstrated that increased activation and synchronization is a trigger of the pathological cascade in this disease. Linked to this last idea, is the other plausible interpretation of sMCI hypersynchronization as a sign of functional network disruption. The fact that all MCI patients as a group showed hypersynchronization as compared to the control group reinforces this interpretation. However, it is a counterintuitive finding that the sMCI group is the one showing highest hypersynchronisation. We interpret it by arguing that the fast converters, our pMCI, showed less heart-evoked brain synchronization because they were in a more advanced stage (i.e., closer to Alzheimer disease), thereby presenting a more disconnected network as shown in AD patients (Stam, 2010). In contrast, sMCI subjects are at an earlier stage and therefore still able to demonstrate a higher response to the heart dynamics. This hypersynchronization, induced by the impaired inhibitory response showed in sMCI patients, may give rise to a vicious cycle increasing the likelihood of brain amyloid deposition (Cirrito et al., 2008). It has to be tested, in humans, whether the reduction of this brain response by anti-epileptic drugs (Sanchez et al., 2012) could delay the typical 15% rate of yearly conversion of MCI into Alzheimer disease. We compared sMCI and pMCI APOE4 carriers. The first group showed again higher heart-evoked brain synchronization, which clearly suggests that these two groups represent two separate stages of the disease showing different heart-evoked brain synchronization. As indicated by the results from the SVM technique, putting the heart into it improves our ability to distinguish between both stages as compared to using brain synchronization alone.

### Implications for cognitive neuroscience

Considering the heartbeat as a temporal trigger to study resting (RS) state unveil interesting phenomena in brain dynamics. In fact, we showed that heart-evoked responses correlated with cognitive performance and dementia status not only in somatosensory areas but also in frontal areas and regions belonging to the DMN (Figure 1). The heart and brain dynamics interact not only in response to the heartbeat but along the cardiac cycle. This continuous connection would be one of the factors responsible for the richness of rhythms observed in brain activity and the aberrant brain network found in pathology (figure 2 and 3). Our results show, for the first time, that considering both heart and brain dynamics simultaneously enriches the study when compared with the cerebral activity alone. As shown in Figure 2B (black bars) results are not reproduced using brain data not locked to heartbeats (activity randomly sampled at intervals mimicking heart pulses) and the average of RS intervals supposes a waste of information. Additionally, based on machine learning results we show that the prediction to progression to AD is higher from the heart-brain interaction than from brain synchronization (Figure 4). It strongly suggests the need to rethink the role of the heart in cognitive neuroscience.

## Acknowledgements and Funding

This work was supported by the Nirakara Institute and by projects from the Spanish Ministry of Economy and Competitiveness (PSI2009-14415-C03-01 and PSI2012-38375-C03-01), and a postdoctoral fellowship to M.E.L. (FJCI-2014-22730). We thank Dr. Ricardo Bajo, and Dr. Juan José G. Galarraga for technical support, useful suggestions and strong inspiration.

